# Exosomal miR-145-5p promotes apoptosis of renal tubule epithelial cells through the JNK signalling pathway

**DOI:** 10.1101/2025.01.08.632043

**Authors:** Xiao-long Deng, Le-ping Yan, Yuan Lan, Jun-miao Guo, Run-qi Yuan, Rong Zhang, Pu Mao

## Abstract

Acute kidney injury (AKI) is the most prevalent extrapulmonary organ failure observed in patients with acute respiratory distress syndrome (ARDS). However, the underlying mechanisms leading to AKI in ARDS remain unclear. Exosomes facilitate intercellular and inter-organ communication via exosomal microRNAs (miRNAs), which potentially contribute to the crosstalk between the lung and kidney in ARDS. Therefore, the purposes of our study were to investigate the involvement of exosomal miRNAs in the development of AKI in ARDS. In current study, Sprague-Dawley rats were induced to develop ARDS via hydrochloric acid aspiration. Nanoparticle tracking analysis, Transmission electron microscopy and western blotting were conducted to analyse the exosomes. Quantitative reverse transcription-polymerase chain reaction and *in situ* hybridisation showed elevated expression level of miR-145-5p in the circulating exosomes and lung tissue of ARDS rats. Overexpression of miR-145-5p disrupted the proteomic profile of renal proximal tubule epithelial cells, leading to impacts on proteins associated with various biological processes, including apoptosis. We further demonstrate that miR-145-5p overexpression enhanced JNK phosphorylation by directly repressing RBM3 expression and promoted cell apoptosis. *In* vivo, administration of agomiR-145-5p injection via intravenous route, which subsequently induced renal tubular cell apoptosis. These findings have revealed that miR-145-5p may serve as a novel mediator for the apoptosis of renal cell in the context of ARDS.

## 1. Introduction

Acute respiratory distress syndrome (ARDS) is a life-threatening lung injury in the intensive care unit (ICU), frequently leads to the development of acute kidney injury (AKI) as a significant and prevalent complication^1,2^. In patients with ARDS admitted to an ICU, the AKI prevalence is 44.3% with mortality rates ranging from 20.2% to 42.3%^3,4^.

The potential mechanisms underlying the development of AKI in patients with ARDS involve acidosis and blood gas disturbances ^5–7^. The systemic release of pro-inflammatory mediators from injured lungs has also been considered a type of biological injury resulting in AKI^8^. Nevertheless, numerous gaps remain regarding the molecular mechanism for AKI development among critically ill patients^9^.

MicroRNAs (miRNAs) are short non-coding RNAs that play a regulatory role in gene expression by suppressing translation or degrading targeted mRNAs, exerting a strong influence on gene regulation^10^. Moreover, miRNAs are integral constituents of exosomes. Those exosomes are small (30-150nm) endogenous membrane vesicles secreted by various cell types and play a crucial role in mediating intercellular communication and crosstalk between organs by delivering varied signal molecules, comprising miRNAs, proteins, and mRNAs et al^11,12^. In ARDS, the miRNA expression levels in circulating exosomes are reportedly altered and exhibits significant diagnostic and prognostic utility^13–15^. These findings suggest that exosomal miRNAs may participate in the inter-organ crosstalk in ARDS.

The pathogenesis of AKI is multifactorial but definitely involves renal cell apoptosis. Proximal tubule epithelial cells are highly susceptible to apoptosis and this process contribute to the development of AKI^16,17^. We posit that altered miRNA expression in circulating exosomes contributes to human renal proximal tubule epithelial cell apoptosis in ARDS, resulting in the development of AKI.

In this study, we used acid aspiration to model ARDS in rats and assess the differential expression of the apoptosis-related miRNAs in plasma exosome. then, we primarily investigated the impact of exosomal mir-145 on the regulation of apoptosis in renal tubule epithelial cells and its contribution to the development of AKI in the context of ARDS.

## 2 Materials and methods

### 2.1 Rat model of ARDS

Male Sprague-Dawley (SD) rats (300-350 g in weight) were procured from the Guangdong Medical Animal Experiment Centre. The rats were anesthetized intraperitoneally using a combination of 10% chloral hydrate and 25% urethane (1:1) at a dosage of 0.5 mL per 100 g body weight. Afterwards, an endotracheal tube was inserted with a 14-G artery indwelling needle cannula. Hydrochloric acid (HCl; pH = 1.0, 1.5 mL/kg) or normal saline (1.5 mL/kg; healthy control) was administered through the endotracheal tube. To ensure a uniform distribution of HCl or saline in the lungs, the rats underwent mechanical ventilation for 10 min with the aid of a Servo-i ventilator (Maquet-Dynamed Inc, Solna, Sweden) with the following ventilation parameters: FiO_2_ = 100%, PEEP = 3 cm H_2_O, PC above PEEP = 13 cm H_2_O, and RR = 60 times per min. After 24 h, 0.1 mL of abdominal aortic blood was extracted for assessing the arterial blood gases (i-STAT (Abbott Laboratories, Abbott Park, IL). Blood samples were retrieved from the inferior vena cava, while the lungs and kidneys were surgically removed and stored at −80 ℃ for subsequent analysis or immersed in 4% paraformaldehyde solution for histological evaluation. The arterial oxygenation index (PaO_2_/FiO_2_ ratio) and pathological changes in the lungs were assessed to verify the establishment of the rat model of ARDS. The total lung injury score was calculated using a previously published protocol^18,19^. Two experienced pathologists evaluated the lung injury scores in a blinded manner, based on the extent of neutrophil infiltration, alveolar collapse, congestion, haemorrhage, macrophage infiltration, as well as alveolar wall thickness or hyaline membrane formation. The severity of the lung injury was measured on a scale of 0 to 3 (normal, mild, moderate, and severe, respectively). This research was subjected to approval by the Animal Ethics Committee of Guangzhou Medical University (No. GY2019-090).

### 2.2 Exosome uptake assay

PKH26 Red Fluorescent Cell Linker Mini Kit (Sigma, MINI26) was employed to label exosomes as per the instructions. The PKH26 dye was diluted in 100 μL of diluent C to attain a final concentration of 2 μM. Resuspended exosomes were introduced into 100 μL of the exosome-Diluent C solution and maintained for 5 min at 37℃. Subsequently, 500 μL of 10% exosome-free FBS was added to bind excess dye. PBS was then added with the final volume brought to 12.5 mL and ultracentrifuged at 100,000 × g for 120 min at 4°C. The labelled exosomes were then resuspended in 50 μL of PBS. NRK-52E cells were cultured in confocal glass-bottomed petri dishes with labelled exosomes and stained with DAPI (Sigma). Confocal microscopy (LSM 880, ZEISS, Oberkochen, Germany) was used to visualize exosome uptake.

### 2.3 Isolation of plasma exosomes

Total Exosome Isolation (from plasma) kit from Invitrogen (Carlsbad, USA), as per the the manufacture’s protocol, was utilized to isolate the plasma exosomes. They were subsequently resuspended in phosphate-buffered saline (PBS) for further analysis and examined using a Hitachi H-7650 transmission electron microscope (Hitachi High-Technologies, Tokyo, Japan). Nanoparticle tracking analysis (NTA) was performed at the Guangzhou NoVa Biological Corporation.

### 2.4 Quantification of exosomal miRNAs

Exosomal RNAs were extracted with the aid of the exoRNeasy MiDi Kit (Qiagen, Hilden, Germany) for Small RNA following the manufacturer’s protocol. For exosomal RNA isolation, cel-miR-39 was added to normalise the technical variation between the samples before the chloroform step. NanoDrop 2000 Spectrophotometer (Thermo Fisher Scientific, Wilmington, DE) was utilized to verify RNA concentrations. The cDNA was synthesised using the miRCURY LNA Reverse Transcription Kit (Qiagen). Briefly, 2 µL of template RNA from each group was mixed with 2 µL of 5× miRCURY SYBR® Green RT Reaction Buffer, 1 µL of 10× miRCURY RT Enzyme Mix, and 4.5 µL of RNase-free water. The mixtures were incubated at 42°C followed by a 5 min heat activation step at 95°C to inactivate the reverse transcriptase, and then promptly cooled to 4°C. Subsequently, real-time PCR utilizing the miRCURY LNA SYBR® Green PCR Kit (Qiagen) was executed. The 10 µL RT reaction was diluted in 290 µL of RNase-free water (1:30); 3 µL of cDNA template was mixed with 5 µL of 2× miRCURY SYBR® Green Master Mix, 1 µL of Resuspended miRNA-specific primer mix, and 1 µL of RNase-free water. The mixtures were subjected to incubation for 2 min at 95°C, followed by 40 cycles for 10 s at 95°C, and 56°C for 60 s. The specific primers and external RNA reference cel-miR-39-3p (5’-UCACCGGGUGUAAAUCAGCUUG-3’) were purchased from Qiagen. The relative expression level of each miRNA was normalized to cel-miR-39-3p and calculated using the 2^-ΔΔ*Ct*^ method.

### 2.5 Cell culture and transfection

The NRK-52E renal tubular epithelial cells and human renal proximal tubule epithelial cells (RPTECs; ATCC CRL-4031^TM^) were accessed at the American Type Culture Collection (ATCC, Manassas, VA). The NRK-52E cells were grown in Dulbecco’s modified eagle medium (DMEM) supplemented with high glucose and glutamine (Gibco, Carlsbad, CA) along with 5% fetal bovine serum (FBS; Gibco). RPTECs were grown in DMEM/F12 (Gibco), along with 1% RPTEC Supplement A (ATCC), 1.6% RPTEC Supplement B (ATCC), and G418 (0.1 mg/mL; Sigma-Aldrich). BEAS-2B cells, healthy human bronchial epithelial cells (ATCC), were cultivated in 10% FBS-contained DMEM/F12. All cells were exposed to 95% air and 5% CO_2_ and incubated at 37°C.

The BEAS 2B cells were grown in 6 well plates at a density of 2 × 10^5^ cells per well until they attained confluence. Subsequently, the cells were kept in a hypoxic environment for 6 or 12 hours, under 1.0% O_2_, 5% CO_2_, and 93.5% N_2_, at 37°C. For the normoxic control group, BEAS-2B cells were cultured under 95% O_2_ and 5% CO_2_ at 37°C.

The miRNA mimic, miRNA inhibitor, RBM3 siRNA, and scramble RNA controls were procured from RiboBio (Guangzhou, China). Scramble RNA transfected cells were utilized as the control group (referred to as “Scramble”), whereas miR-145-5p mimic and inhibitor (30 nM) were utilized to transfect the RPTECs with the aid of the Lipofectamine™ RNAiMAX Transfection Reagent (Thermo Fisher Scientific, Waltham, MA). The sequence of the miR-145-5p mimics was 5’-GUCCAGUUUUCCCAGGAAUCCCU-3’, whereas that of the miR-145-5p inhibitor was 5’-AGGGAUUCCUGGGAAAACUGGAC-3’. The sequence of si-RBM3 was as follows: sense strand, 5’-UGGCAGGUAUUAUGACAGUdTdT-3’, anti-sense strand, 5’-ACUGUCAUAAUACCUGCCAdTdT-3’.

### 2.6 Data-independent acquisition (DIA) proteomic analysis

RPTECs were transfected with miR-145-5p mimic (30 nM) or scramble RNA controls (30 nM) for 48 h; Subsequently, the cells were harvested to extract proteins, which were further subjected to digestion using sequence-grade modified trypsin. For DDA library construction and quality control, the resulting peptide mixture was separated using XBridge C18 column (Waters, Milford, MA) equipped with the high-performance liquid chromatography system Ultimate 3000 (Thermo Fisher Scientific). The fractions were then injected into a Thermo Fisher Scientific Orbitrap Lumos HF mass spectrometer with a connected EASY-nLC 1200 chromatography system (Thermo Fisher Scientific) for DDA library generation. DIA data-independent acquisition was employed to perform subsequent quantitative proteomic analysis. The mass spectrum parameters were configured as follows: MS scanning range (m/z) of 350-1500, DIA scan resolution of 120,000, automatic gain control (AGC) target set at 4e6, maximum injection time of 50 ms, HCD-MS/MS resolution of 30,000, AGC target set at 1e6, collision energy of 32, energy increase of 5%, variable window acquisition with 60 windows, and each window overlapping by 1 m/z. The resulting data were then assessed with the aid of the Spectronaut X software (Biognosys AG, Schlieren, Switzerland) and examined using the Universal Protein database. Proteins exhibiting a fold change of > 1.5 or < 0.67 and *P* value < 0.05 were deemed as differentially expressed proteins (DEPs). The DEPs were subjected to Gene ontology (GO) and Kyoto Encyclopaedia of Genes and Genomes (KEGG) analysis to perform functional annotation and pathway enrichment analysis. Gene set enrichment analysis (GSEA) was employed to evaluate both negatively and positively correlated DEPs. The protein sequencing procedures and bioinformatics analysis were conducted by Gene Denovo Bio-tech Co. (Guangzhou, China). The proteomics and raw data from this study have been submitted to iProX under accession number IPX0006160000.

### 2.7 Western blotting

The samples were lysed using RIPA Lysis Buffer and phenylmethanesulfonylfluoride (PMSF) at a final concentration of 1 mM, whereas the protein levels were examined through Bicinchoninic acid protein assay kit (Thermo Scientific, Ottawa, Canada). Separation of the protein was conducted through sodium dodecyl sulphate– polyacrylamide gel electrophoresis (SDS-PAGE) for 30 mins at 80 V, followed by 1 h at 100 V. These were then shifted onto polyvinylidene difluoride membranes (Bio-Rad, Richmond, CA) and maintained at 200 mA for 2 h. To block the membranes, 5% non-fat milk in TBST was added for 1 h, along with subsequent overnight incubation with primary antibodies against various proteins at 4°C. These proteins included TSG101 (sc-7964, 1:1000), p-p38(D-8) (sc-7973, 1:1500), p-JNK (G-7) (sc-6254, 1:1500), JNK (sc-7345, 1:1500), p38 (sc-7972, 1:1500), AIP1 (sc-365921, 1:1500), Calnexin (sc-23954, 1:1000) (Santa Cruz Biotechnology, Santa Cruz, CA); PARP (9542T, 1:1000), Bax (2772S, 1:1000), Bcl-2 (4223S, 1:1000) (Cell Signalling Technology, Danvers, MA); CD81 (ab109201, 1:2000), Bak (ab32371, 1:5000), RBM3 (ab134946, 1:3000), Caspase-3 (ab32351, 1:5000) (Abcam, Cambridge, MA). Afterwards, the membranes were exposed to the relevant HRP-linked secondary anti-rabbit IgG (7076S, 1:3000) and anti-mouse IgG (7074S, 1:3000) (Cell Signalling Technology) antibodies for 1 h at 25°C. Visualization of the protein bands was achieved through an enhanced chemiluminescence scanner (PXi9, Syngene, Frederick, MD).

### 2.8 Luciferase reporter assay

The pGL3-RBM3 3′UTR vectors, containing wild-type or mutant-type RBM3 3′UTR, were constructed by GenePharma (Shanghai, China). RPTECs were co-transfected with 100 ng of pGL3-RBM3 3′UTR WT (wild type RBM3 3′UTR) or pGL3-RBM3 3′UTR Mut (mutant-type RBM3 3′UTR) as well as 30 nM miR-145-5p mimic or scramble control with the aid Lipofectamine-2000 (Invitrogen) as per relevant protocols. Luciferase reporter assay was performed following cell collection (after 48 h) using the Dual-Luciferase Reporter Assay System (Promega, Madison, WI).

### 2.9 Flow cytometry

Propidium iodide (PI) and Fluorescein isothiocyanate (FITC)-conjugated Annexin V (Abcam) were employed to detect apoptotic cells. Briefly, cells were digested with StemPro® Accutase® Cell Dissociation Reagent (Gibco), collected in centrifuge tubes, and centrifuged at 4°C for 5 min at 300 *× g*. The supernatant was discarded, and the cells were rinsed thrice with PBS (5 min centrifugation at 300 *× g* at 4°C); this step was repeated thrice. Resuspension of the cells in 1× binding buffer was carried out after discarding the PBS. Cells were subjected to Annexin V and PI staining solution and incubated for 10 min at room temperature away from light. Data were retrieved using a BD FACSCantoII (BD Biosciences, Heidelberg, Germany) and analysed with FlowJo software (Tree Star Inc., Ashland, OR).

### 2.10 JNK Inhibitor Experiment

RPTECs were exposed to a JNK inhibitor SP600125 (Beyotime Biotechnology, Shanghai, China) at 20 μM for 2 h. And then the miR-145-5p mimic, RBM3 siRNA, or scramble RNA controls (30 nM all) were transfected into the RPTECs. DMSO served as a control.

### 2.11 agomiRNA injection

SD rats were adaptively fed for 5 days and randomly classified into two groups (*n* = 5 mice/group minimum). The agomiRNA-145-5p and agomiR negative control were purchased from RiboBio (Guangzhou, China). Briefly, 50 nmol agomiR-145-5p and agomiR negative control were diluted with 200 μL of normal saline and placed on ice. The rats were administered with the mixtures via tail vein injection. After 48 h, the kidneys were excised and stored at −80℃ for further analyses or soaked in a 4% paraformaldehyde solution for histological evaluation.

### 2.12 miRNA *in situ* hybridisation

RNAscope^TM^ Assay and the miRNAscope™ HD Detection Kit-RED (Advanced Cell Diagnostics, Newark, NJ) was employed to detected miR-145-5p in lung and kidney tissue as described in the instruments^20^. The formalin fixed and embedded in paraffin tissue sections were subjected to deparaffinization and rehydration using xylene, followed by 100% ethanol. the sections were pre-treated with RNAscope^TM^ Assay (Advanced Cell Diagnostics, Newark, NJ). Briefly, the sections were incubated in 10% neutral buffered formalin at room temperature overnight. The deparaffinised samples were then treated for 10 min with hydrogen peroxide at room temperature. The sections were subjected to boiling for 15 min in 1X Target Retrieval Reagents solution and subsequently covered with Protease III in a pre-warmed HybEZ™ Humidity Control Tray at 40°C. Afterwards, the sections were subjected to the miRNAscope™ HD Detection Kit-RED (Advanced Cell Diagnostics). After removing excess liquid from the sections, the slides were covered with miRNAscope™ miR-145-5p, RNU6, or scramble probe and subjected to incubation at 40°C for 2 h. Hybridize Amp1 to 6 were sequentially added to the sections and incubated. Subsequently, the sections were stained for miR-145-5p with a 1:60 ratio of Fast Red -B to Fast Red -A and subjected to counterstaining with 50% hematoxylin staining solution and 0.02% ammonia water. After washing and drying the sections, they were sealed with Vectamont mounting medium (Vector Laboratories, Inc., Newark, CA) for further evaluation of the target probe signal. The signals were visually scored using a scale ranging from 0-3, with respective notations as follows: 0, no staining or < 1 dot/cell at 40× magnification; 1, 2–10 dots/cell, with no or negligible cell clusters at 40× magnification; 2, 11–20 dots/cell and/or < 25% dots in clusters at 20–40× magnification; 3, > 20 dots per cell and/or > 25% of dots in clusters at 20× magnification.

### 2.13 Histological and immunohistochemical staining

Lung tissues were subjected to staining by means of haematoxylin and eosin (H&E), as previously outlined (1). The kidney tissue samples were subjected to deparaffinization using xylene, followed by rehydration with xylene with graded ethanol. Boiling sodium citrate-hydrochloric acid buffer solution was used to immerse the sections, which were heated in the microwave for 5 min, repeating the process thrice, to achieve antigen retrieval. After the samples were cooled to room temperature, endogenous peroxides were removed with 3% hydrogen peroxides and washed with PBS thrice. To block the sections, 5% bovine serum albumin was utilized for 30 min at 37°C. Afterward, incubation with anti-cleaved caspase-3 antibodies (9664, 1:200) (Cell Signalling Technology) was conducted at 4°C for a whole night. The samples were subjected to incubation with secondary antibodies after washing them with PBS thrice. Following three additional washes, diaminobenzidine was applied for approximately 5 minutes, and haematoxylin was used as a counterstain for 2 minutes. Subsequently, ethanol was used to dehydrate, xylene was used to vitrify, and neutral resin was employed to seal the sections.

### 2.14 TUNEL staining

Renal apoptosis was assessed with the TUNEL Apoptosis Detection Kit (Servicebio technology co., Wuhan, China). Briefly, fixed-frozen renal tissue sections were fixed with 4% paraformaldehyde solution and exposed at 37°C to Proteinase K (20 μg/mL) for 25 min. Next, the samples were treated with the TUNEL reaction mixture comprising of FITC-12-dUTP Labelling Mix, and TUNEL-positive nuclei were visualised using fluorescence microscopy.

### 2.15 Statistical analysis

All data were presented as mean ± SEM of *n* independent experiments. GraphPad Prism 9 software (GraphPad Software Inc., San Diego, CA) was employed to conduct statistical analyses, utilizing a student’s *t*-test for the comparative assessment of two groups, whereas one-way analysis of variance was performed for the comparative assessment of multiple groups. Statistically, a *P* value of < 0.05 indicated the significance level. Independent experiments were repeated thrice.

## 3 Result

### 3.1 Establishment of the rat model of ARDS

To verify the successful establishment of ARDS, H&E staining was performed in lung tissue, and the arterial oxygenation index (PaO_2_/FIO_2_) was assessed. Following 24 hours of HCl instillation, lung tissues obtained from HCl-induced rats exhibited considerable pathological alterations, including interstitial alveolar collapse, alveolar hyperaemia, and inflammatory cell infiltration, as compared to lung tissues from the control group rats, as evidenced by Figure 1A and 1B. Moreover, the PaO_2_/FiO_2_ ratio was significantly decreased in HCl group than the control group (Figure 1C).

**Figure. 1.**
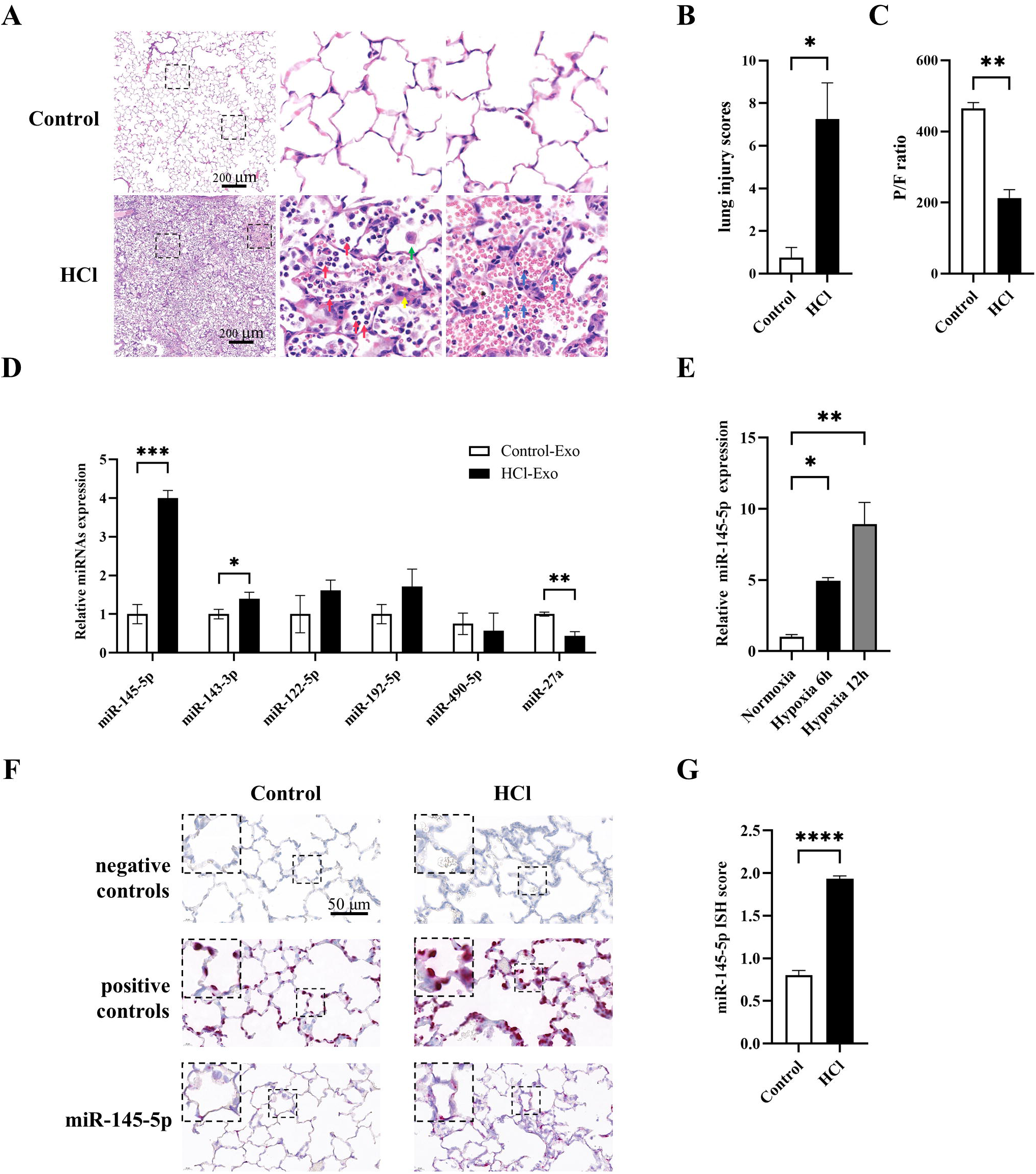
Identification of exosome-derived miRNAs in ARDS. **(A)** H&E staining of lung tissue sections from rats treated with hydrochloric acid or normal saline instillation. (Red arrow, alveolar polymorphonuclear leukocytes; Blue arrow, alveolar haemorrhage; Yellow arrow, alveolar edema, Green arrow, alveolar macrophages; Scale bar 200). **(B)** the quantitative analysis of lung histological damage. **(C)** The arterial oxygenation index was calculated as PaO_2_/FiO_2_. *n* = 5 per group. **(D)** miRNA expression in the plasma exosomes from rats subjected to normal saline or HCl aspiration. *n* = 5 per group. **(E)** miR-145-5p expression in BEAS-2B cells following hypoxia exposure for 6 h or 12 h. **(F and G)** Representative images of miRNA in situ hybridisation in paraffin-embedded lung tissue. RNU6 was used as a positive control; probe-SR-Scramble served as a negative control. The presence of miRNA-145 is shown by red punctate dots in the cytoplasm. The semi-quantitative ISH score was calculated as per the counted number of punctate dots within each cell boundary. Scale bar 50 μm. \**P* < 0.05; \*\**P* < 0.01; \*\*\**P* < 0.001; \*\*\*\**P* < 0.0001.

### 3.2 Characterisation of exosomes

To evaluate the characteristics of exosomes isolated from rat plasma, transmission electron microscopy (TEM) was utilized to observe their size and shape. TEM analysis of isolated fractions revealed typical bilayer membrane vesicles with diameters ranging from 40 to 150 nm as depicted in Figure 2A. Furthermore, the appearance of exosome marker proteins (TSG101 and CD81; Figure 2B) was confirmed by Western blot analysis. Moreover, nanoparticle tracking analysis revealed an exosome concentration of approximately 3.3 × 10^10^ particles/mL. The mean particle size was 135.5nm with a peak at 137 nm (Figure 2C).

**Figure. 2.**
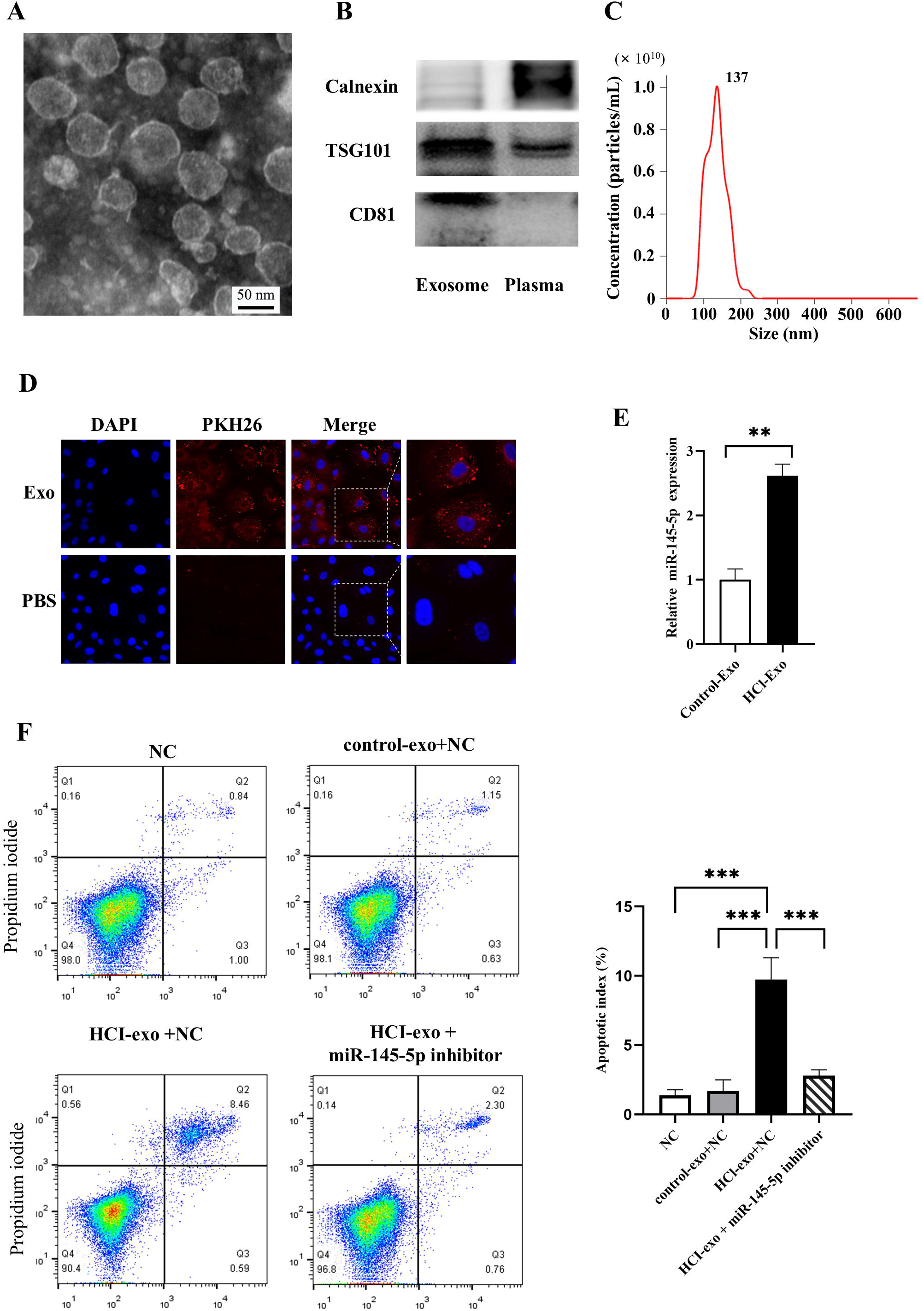
Exosome characteristics and uptake. **(A)** Representative transmission electron micrograph images of the exosomes that were isolated. Scale bar 50 nm. **(B)** Representative blots of TSG101 and CD81 marker proteins in exosomes. **(C)** Representative results of nanoparticle tracking analysis demonstrating size distribution of exosomes. **(D)** Culture of NRK-52E cells in the presence or lack (control) of PKH26-labelled exosomes (red) for 12 h at 37 °C. Nuclei were labelled with DAPI (blue). Scale bar 50 μm. **(E)** miR-145-5p expression in NRK-52E cells treated with plasma exosomes from HCl and control group. **(F)** Flow cytometric analysis with Annexin V-PI staining in NRK-52E incubated with an inhibitor negative control (NC), control-exo+NC, HCI-exo+NC, HCI-exo+miR-145-5p inhibitor for 48 h (*n* = 3 independent experiments). \*\*\*\**P* < 0.0001.

### 3.3 Transfer of miR-145-5p through exosomes and induces apoptosis

The rat renal proximal tubular cell NRK-52E were incubated with PKH26-labelled exosomes, and confocal microscopy revealed that exosomes were internalised by NRK-52E cells after 12 hours of incubation, as depicted in Figure 2D. The uptake of miR-145-5p by NRK-52E cells was then determined by qRT-PCR. We found a 2-fold increase of intracellular miR-145-5p in NRK-52E cells incubated with exosomes derived from HCl group compared to healthy controls (Fig. 2E). These findings provide evidence that miR-145-5p can be transferred to NRK-52E cells through exosomes. Next flow cytometry was conducted to exam the impact on apoptosis after the NRK-52E incubated with exosomes. Exosomes purified from plasma of HCl-induced ARDS rats were found to induced apoptosis. This effect was attenuated by miR-145-5p inhibitor (Fig. 2F).

### 3.4 miRNA expression in plasma exosomes isolated from ARDS rat model

To explore the differences in the miRNAs within exosomes and their potential role in aggravating renal injury, apoptosis-related miRNAs (i.e., miR-145-5p, miR-143-3p, miR-122-5p, miR-192-5p, miR-490-5p, miR-27a) were selected according to the Diana tools database GO:0043065 and analysed their expression using qRT-PCR (Figure 1D). Among the six miRNAs, miR-145-5p and miR-143-3p levels were elevated, while the miR-27a level was reduced in HCL-Exo in comparison with Control-Exo (Figure 1D). More specifically, miR-145-5p expression increased by 4-fold in the HCL-Exo groups (Figure 1D). Considering that miR-145-5p is conserved across species (Appendix Figure 1), it was selected for subsequent functional validation.

To elucidate the potential sources of increased plasma exosomal miR-145-5p in the ARDS model, *in situ* hybridization was conducted for miR-145-5p localization within the lungs of various groups. The smooth muscle layer of vessels and bronchi depicted positive staining for miR-145-5p in the lungs (Appendix Figure 2A), which was consistent with a previous report^21^. Notably, *in situ* staining also revealed the elevated expression of miR-145-5p in the pulmonary epithelial cells after HCI simulation (Figure 1F and 1G). Furthermore, the impact of hypoxia on miR-145-5p production was examined by exposing human lung epithelial cells (BEAS-2B) under hypoxic conditions for 6 or 12 hours. After 12 h, the expression of miR-145-5p was significantly increased in hypoxic compared with normoxia (Figure 1E).

### 3.5 RBM3 was the targets of miR-145-5p

To analyse the impact of miR-145-5p on functional pathways and molecules, data-independent acquisition (DIA) proteomics was conducted on RPTECs transfected with either miR-145-5p mimic or scramble control. A total of 161 differentially expressed proteins (DEPs) were detected (Table S1), of which 131 were upregulated and 30 were downregulated compared with the scramble control (Figure 3A). The proteomics and raw data from this study have been submitted to iProX under accession number IPX0006160000.Gene set enrichment analysis (GSEA) of the DEPs showed that the expression miR-145-5p activated apoptosis and Jak-STAT signalling pathways (Figure 3B). Moreover, proteins involved in cell cycling and DNA replication signalling pathways were remarkably inhibited (false discovery rate (FDR) < 0.25; normalised enrichment score (NES) > 1, Figure 3B). According to recent molecular mechanism research, most miRNAs regulate gene expression via post-transcriptional repression^10^. Thus, we focused on downregulated DEPs and combined the results from two bioinformatic programs of TargetScan and miRDB database to predict miR-145-5p targets (Figure 3C). Consequently, RBM3 was recognised as a candidate target of miR-145-5p. Additionally, miR-145-5p binding site on RBM3 is highly conserved in species (Appendix Figure 1).

**Figure. 3.**
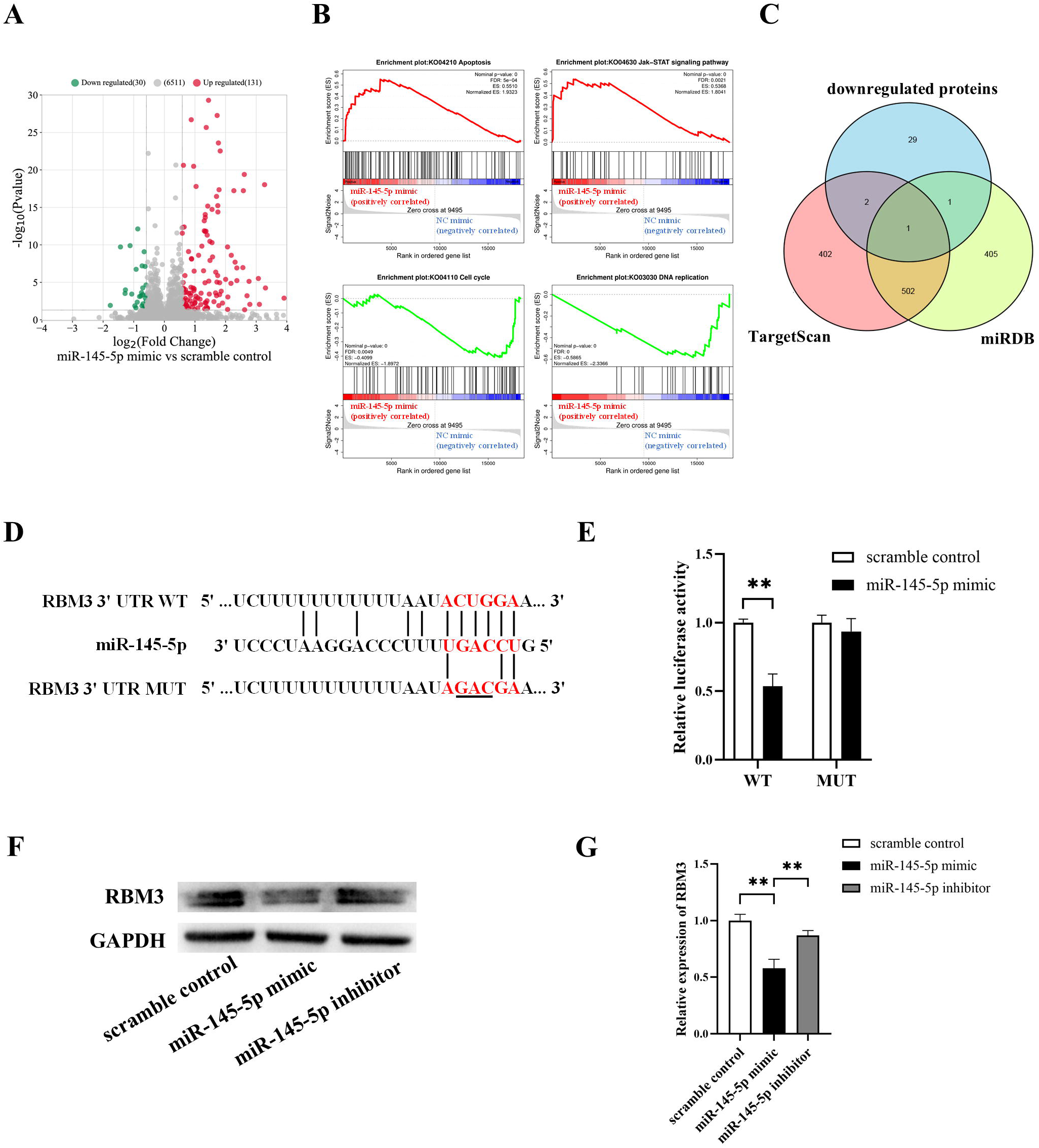
RBM3 is a target gene of miR-145-5p in RPTECs. **(A)** Volcano plot of differentially expressed proteins (DEPs; *P* < 0.05, absolute fold change ≥ 1.5) in miR-145-5p mimic-transfected RPTECs in contrast with the scramble control for 48 h. (**B**) Gene set enrichment analysis (GSEA) of DEPs associated with miR-145-5p. Genes are ranked by signal/noise ratio according to their differential expression between RPTECs transfected with miR-145-5p mimic and scramble control for 48 h; NES, normalized enrichment score. **(C)** Venn diagram of the target prediction for miR-145-5p utilizing the TargetScan, miRDB database, and quantitative proteomic approaches. Quantitative proteomic analysis of proteins downregulated in the miR-145-5p mimic-transfected RPTECs. **(D)** Bioinformatics analysis of the miR-145-5p target sequence in RBM3 3′UTR. The underlined sequence representing the mutant seed sequence in RBM3 3′UTR. **(E)** Luciferase assays of RPTECs co-transfected with miR-145-5p mimic or scramble control and pGL3-RBM3-3′UTR WT (wild type RBM3 3′UTR) or pGL3-RBM3-3′UTR Mut (mutant-type RBM3 3′UTR). **(F and G)** RBM3 protein abundance in RPTECs transfected with miR-145-5p mimic, miR-145-5p inhibitor, or scramble control (n = 4 independent experiments). \*\**P* < 0.01.

To validate that RBM3 is a target of miR-145-5p, pGL3-RBM3 3′UTR WT and pGL3-RBM3 3 ′ UTR Mut were constructed, and a Luciferase reporter assay was performed in RPTECs. Luciferase activity decreased significantly compared with the scrambled control when the miR-145-5p mimic was transfected into pGL3-RBM3-WT. Furthermore, no remarkable difference was observed in luciferase activity following transfection of miR-145-5p mimic with pGL3-RBM3-Mut compared with the scramble control (Figure 3E). Additionally, treatment with miR-145-5p mimic or inhibitor remarkably decreased or elevated the abundance of RBM3 protein, respectively (Figure 3F and 3G). These findings collectively imply that RBM3 is a direct target of miR-145-5p.

### 3.6 miR-145-5p promotes apoptosis in RPTECs

RMB3 has a well-established anti-apoptosis role. For further investigating the association of miR-145-5p with apoptosis, western blotting and flow cytometry were conducted to examine its impact on apoptosis after transfecting RPTECs with miR-145-5p mimic. The resulting data revealed a considerable elevation in the protein levels of PARP, Bak, Bax, and cleaved caspase-3 after miR-145-5p mimic transfection, in contrast with the scramble control (Figure 4A and 4B). Following miR-145-5p mimic transfection, the proportion of late apoptotic cells (Annexin-V^+^/PI^+^) remarkably increased from 1.87% to 18.41% in contrast with the scramble control (Figure 4C). These data provide evidence that miR-145-5p promotes apoptosis in RPTECs.

**Figure. 4.**
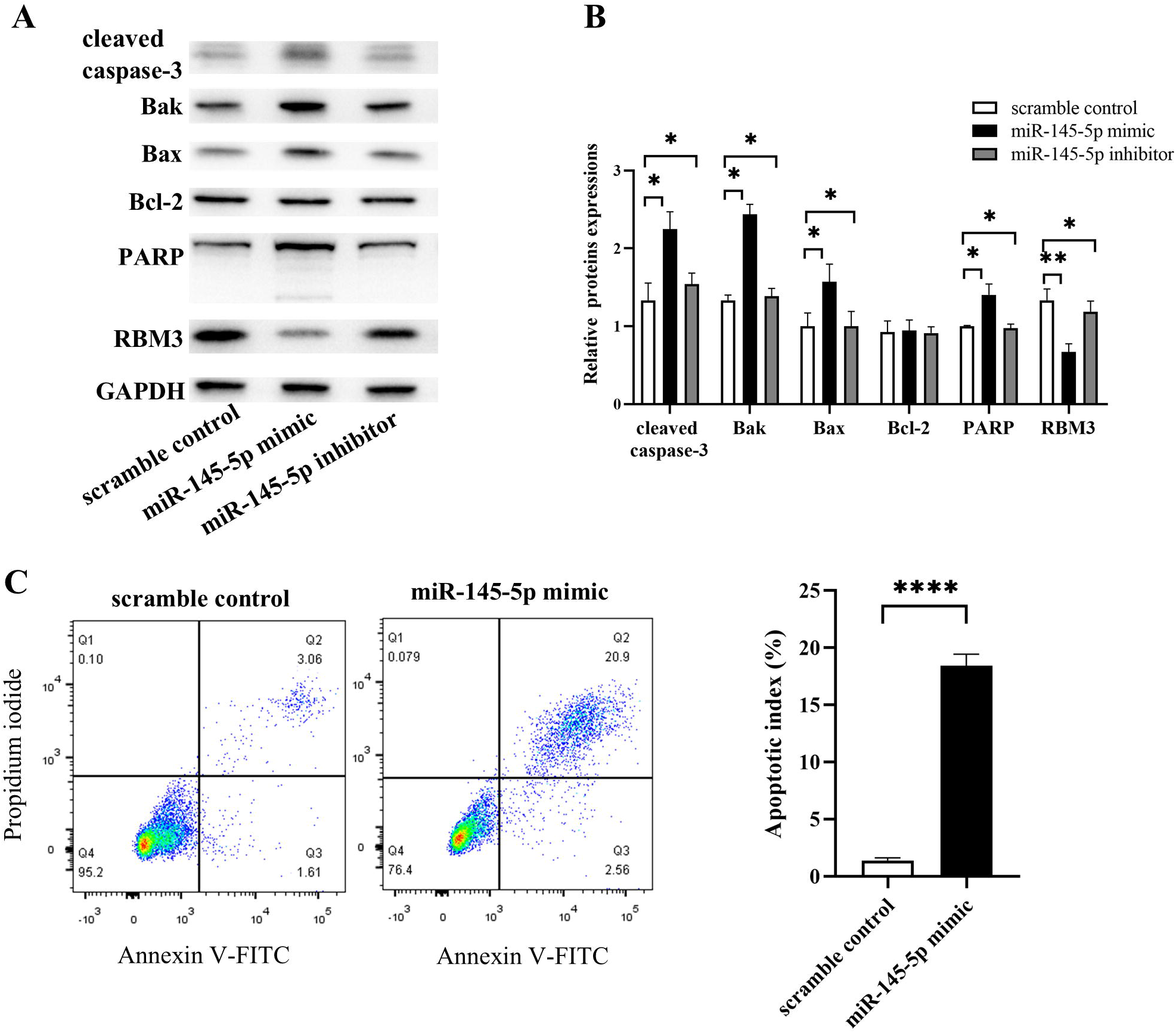
Ectopically expressed miR-145-5p or the transfection of miR-145-5p inhibitor induces apoptosis. **(A and B)** Representative blots and quantified data show apoptosis-related protein expressions in RPTECs transfected with miR-145-5p mimic, miR-145-5p inhibitor, or scramble control. **(C)** Flow cytometric analysis with Annexin V-PI staining in RPTECs transfected with miR-145-5p mimic, or scramble control for 48 h (*n* = 3 independent experiments). \**P* < 0.05, \*\**P* < 0.01 and \*\*\*\**P* < 0.0001.

### 3.7 miR-145-5p enhances JNK phosphorylation in RPTECs by targeting RBM3

The activation of JNK pathway has been shown to promote apoptosis^22,23^. AIP is a key protein contributes to the activation of JNK signalling ^24,25^, which was found increased in the miR-145-5p mimic transfection group compared with the scramble control in RPTECs by the proteomics analysis. Therefore, to elucidate the downstream signalling pathway responsible for miR-145-5p-induced apoptosis in RPTECs, the abundance of JNK pathway proteins was examined. JNK phosphorylation was elevated in RPTECs following treatment with miR-145-5p mimic or RBM3 siRNA (Figure 5A and 5B); whereas p38 phosphorylation was not significantly impacted (Figure 5A and 5B). Correspondingly, the protein levels of AIP1 increased, and RBM3 decreased following treatment with the miR-145-5p mimic or RBM3 siRNA (Figure 5A and 5B). Treatment with the JNK inhibitor (SP600125) attenuated the apoptosis induced by the miR-145-5p mimic or RBM3 siRNA (Figure 5C and 5D). Thus, the outcomes imply that the JNK signalling pathway is the downstream pathway accountable for the apoptosis induced by miR-145-5p in RPTECs.

**Figure. 5.**
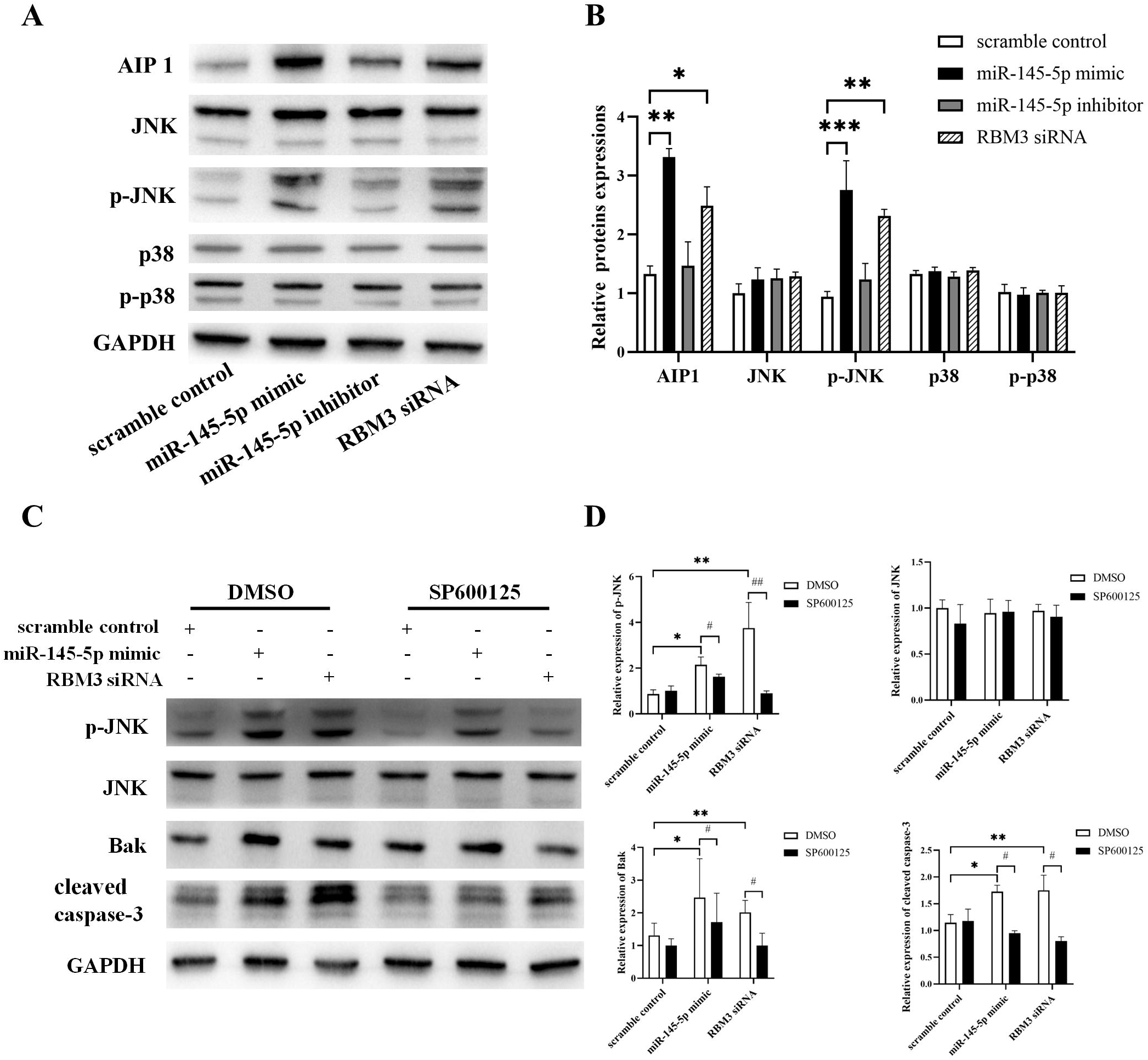
miR-145-5p/RBM3 impact apoptosis through JNK signalling. **(A and B)** Protein expression of AIP 1, p-JNK, JNK, p38, and p-p38 in RPTECs in response to transfection with miR-145-5p, RBM3 siRNA, or scramble control for 48 h. **(C and D)** Protein expression of p-JNK, JNK, Bak, and cleaved caspase-3 in RPTECs in response to treatment with a JNK inhibitor (SP600125) or DMSO for 2 h, followed by transfection with miR-145-5p mimic, RBM3 siRNA, or scramble control. \**P* < 0.05, \*\**P* < 0.01, \*\*\**P* < 0.001 comparison of all groups in DMSO pre-treatment. ^#^*P* < 0.05, ^##^*P* < 0.01 # DMSO pre-treatment versus SP600125 pre-treatment.

### 3.8 AgomiR-145-5p induces renal proximal tubule epithelial cell apoptosis *in vivo*

To verify the function of miR145-5p *in vivo*, we administrated an agomiR-145-5p to rats via the tail vein, then employed ISH to localise miR-145-5p in the rats. Compared with saline or agomiR-NC injection, the miR-145-5p signal (pink area) was enhanced in the renal tubular cells of rats injected with agomiR-145-5p (Figure 6A and 6B). A strong miR-145-5p staining signal was also observed in the smooth muscle layer of vessels in the kidneys (Appendix Figure 2B). Images for negative and positive controls are presented in Appendix Figure 2B. Moreover, the number of TUNEL-positive cells was increased in renal tissue following miR-145-5p administration (Figure 6C and 6D). Meanwhile, the cleaved caspase-3 stain level was increased in the renal tubule epithelial cells following the administration of agomiR-145-5p (Figure 6E). Overall, these data demonstrated that elevation of miR-145-5p enhanced the apoptosis of renal tubule epithelial cells *in vivo*.

**Figure. 6.**
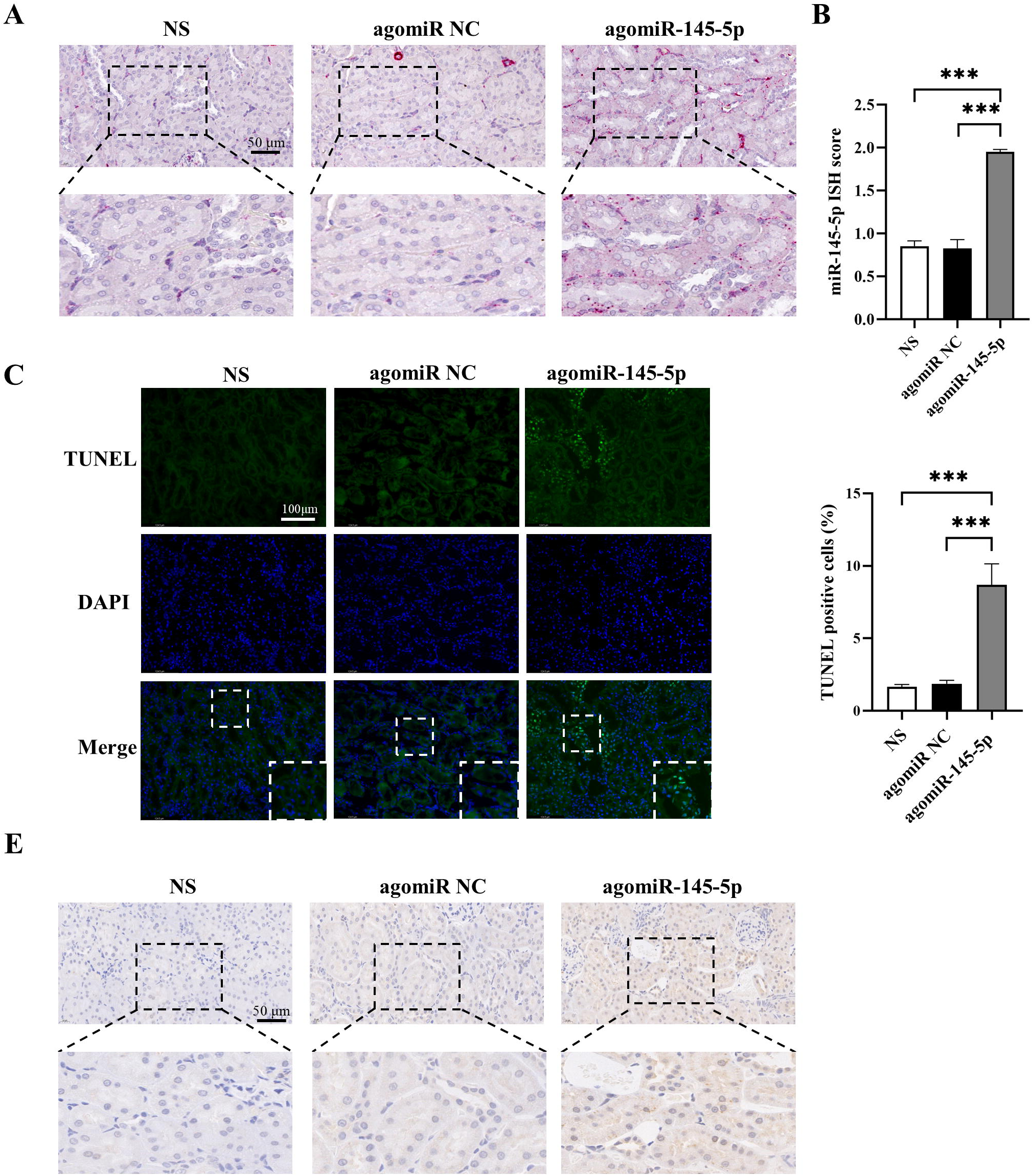
miR-145-5p promotes renal tubule epithelial cell apoptosis *in vivo*. **(A and B)** miR-145-5p signalling (pink-red areas) in rats injected with agomiR-145-5p after 48 h (*n* = 5). Scale bar 50 μm. NS, normal saline. **(C and D)** TUNEL staining for apoptosis in renal tissues after treatment with agomiR-145-5p (*n* = 5). Scale bar 100 μm. **(E)** Representative images of cleaved caspase-3 immunohistochemical staining in kidney tissues. Scale bar 50 μm. \*\*\**P* < 0.001.

## 4 Discussion

In our research, we conducted an assessment of circulating exosomal miRNAs in rats with ARDS to investigate the potential mechanism underlying the development of AKI in ARDS. Our study yielded three primary findings: first, we observed an upregulation of miR-145-5p in plasma exosomes of ARDS rat model; second, RMB3 was demonstrated as the direct target of miR-145-5p; third, miR-145-5p-RBM3-JNK signalling pathway was determined as crucial role in promoting apoptosis of RPTECs. In recent decades, various miRNAs have been found to contribute to ARDS pathogenesis. For instance, Tang *et al*. reported that circulating exosomes from LPS-induced ARDS mice trigger the endoplasmic reticulum to promote ARDS development^26^. Jiang *et al.* discovered that the serum exosome concentration was remarkably increased in acute lung injury mice and exosomal miR-155 contributed to the proliferation and inflammation of macrophages^27^. Meanwhile, Fan *et al*. described that the levels of circulating miR-92a were elevated in patients with sepsis-induced ARDS and contributed to cellular apoptosis and migration *in vitro*^15^. Still further, Parzibut *et al*. identified 12 differentially expressed miRNAs in plasma-derived exosomes from patients with ARDS^13^. Recent discoveries have shed light on the significance of exosome-mediated communication between cells and organs ^28^. This reminiscent of the crosstalk of lung and kidney in the development of ARDS.

Moreover, considering that the instillation of gastric contents is a common risk factor for ARDS^1^ and the detrimental role of apoptosis in RETECs has been demonstrated in multiple models of AKI^29,30^, in our study an ARDS rat model was established through the HCl aspiration-induced lung injury to explore the effect of miRNA in circulation exosome on apoptosis of proximal tubule epithelial cells. Then the circulating exosomes of these rats were analysed, and three apoptosis-associated miRNAs were found to be differentially expressed. The miR-145-5p expression demonstrated the most significant alteration.

miR-145-5p is primarily localized in the smooth muscle layer of pulmonary vessels, bronchi, ureter^21,31^. In our results, *in situ* hybridization localized miR-145-5p within pulmonary epithelial cells as well as the smooth muscle layer of vessels and bronchi when examining the lungs of various groups. The expression of miR-145-5p was remarkably upregulated in the pulmonary epithelial cells of ARDS rats as compared to the control group. Alveolar epithelial cells are known to be the primary source of an extracellular vesicle under non-infectious stimulation^12^, the data from our study suggests that pulmonary epithelial cells may contribute to the increase in plasma exosomal miR-145-5p in ARDS.

Considering that hypoxia is a defining physiological feature in ARDS and that hypoxic response elements (HREs) are associated with the expression of the miR-143/145 promoter region ^32^, BEAS-2B cells were stimulated via hypoxia. Consequently, a sustained and significant upregulation of miR-145-5p was observed in hypoxic BEAS-2B cells. miR-145 expression is also enhanced in the lung tissues of mice exposed to hypoxia^21^. Hence, hypoxia might be responsible for upregulating exosomal miR-145.

RNA-binding motif protein 3 (RBM3) is an outstanding cold shock protein that exhibits essential anti-apoptosis functions^33^. Ma *et al.* reported that RBM3 inhibits the apoptotic response to radiotherapy through the PI3K/AKT/Bcl-2 signalling pathway ^34^. Meanwhile, Feng *et al.* discovered that RBM3 was upregulated in the lung tissue of acute lung injury mice and that its overexpression reduces LPS-induced cell apoptosis ^35^. In our study, RBM3 was determined to be a primary target of miR-145-5p. Furthermore, RBM3 siRNA transfection resulted in increased levels of RPTEC apoptosis and JNK phosphorylation. Meanwhile, inhibition of JNK phosphorylation partially reversed apoptosis in RPTECs. The findings imply that miR-145-5p reduces the abundance of RBM3 and activates the JNK signalling pathway, thus promoting RPTEC apoptosis. The proteomic analysis also confirmed that miR-145-5p overexpression in RPTECs dysregulated the proteomic profile, affecting apoptosis signalling pathways.

This study has some limitations. Firstly, due to limited exosome obtained from rat samples, functional analysis was conducted by transfecting RPTECs with a miR-145-5p mimic. Secondly, the mechanisms related to the storage of miRNAs in exosomes are not fully understood, and further investigation is needed to elucidate precise origins of miR-145-5p.

In conclusion, we demonstrated the increased expression of miR-145-5p in plasma exosomes in rat model of ARDS. We found that miR-145-5p leads to an increase in JNK phosphorylation through the direct repression of RBM3 expression, ultimately resulting in the promotion of cell apoptosis (Figure 7). These findings have revealed a novel mechanism for the apoptosis of renal cell in the context of ARDS.

## Ethical Statement

This study has been approval by the Animal Ethics Committee of Guangzhou Medical University (No. GY2019-090).

## Data Availability statement

Raw data from Figures 1 to 6 were deposited on Mendeley at https://data.mendeley.com/datasets/3xts2ms935/1

**Appendix Table 1.** 161 differentially expressed proteins in RPTECs transfected with miR-145-5p mimic compared with scramble control.

**Appendix Figure. 1** Conservation of the miR-145-5p target sequence in RBM3 3′UTR among different species and conservation of the miR-145-5p sequence among different species.

**Appendix Figure. 2 (A)** In situ hybridisation of miR-145-5p within the smooth muscle layer of vessels and bronchi in the lungs of rats. Scale bar 50 μm. **(B)** Representative images of miRNA in situ hybridisation for RNU6, probe-SR-Scramble, and miR-145-5p in paraffin-embedded kidney tissue. RNU6 was used as a positive control and probe-SR-Scramble served as a negative control. Positive control signals appear as red dots in cells. Scale bar 50 μm.

---

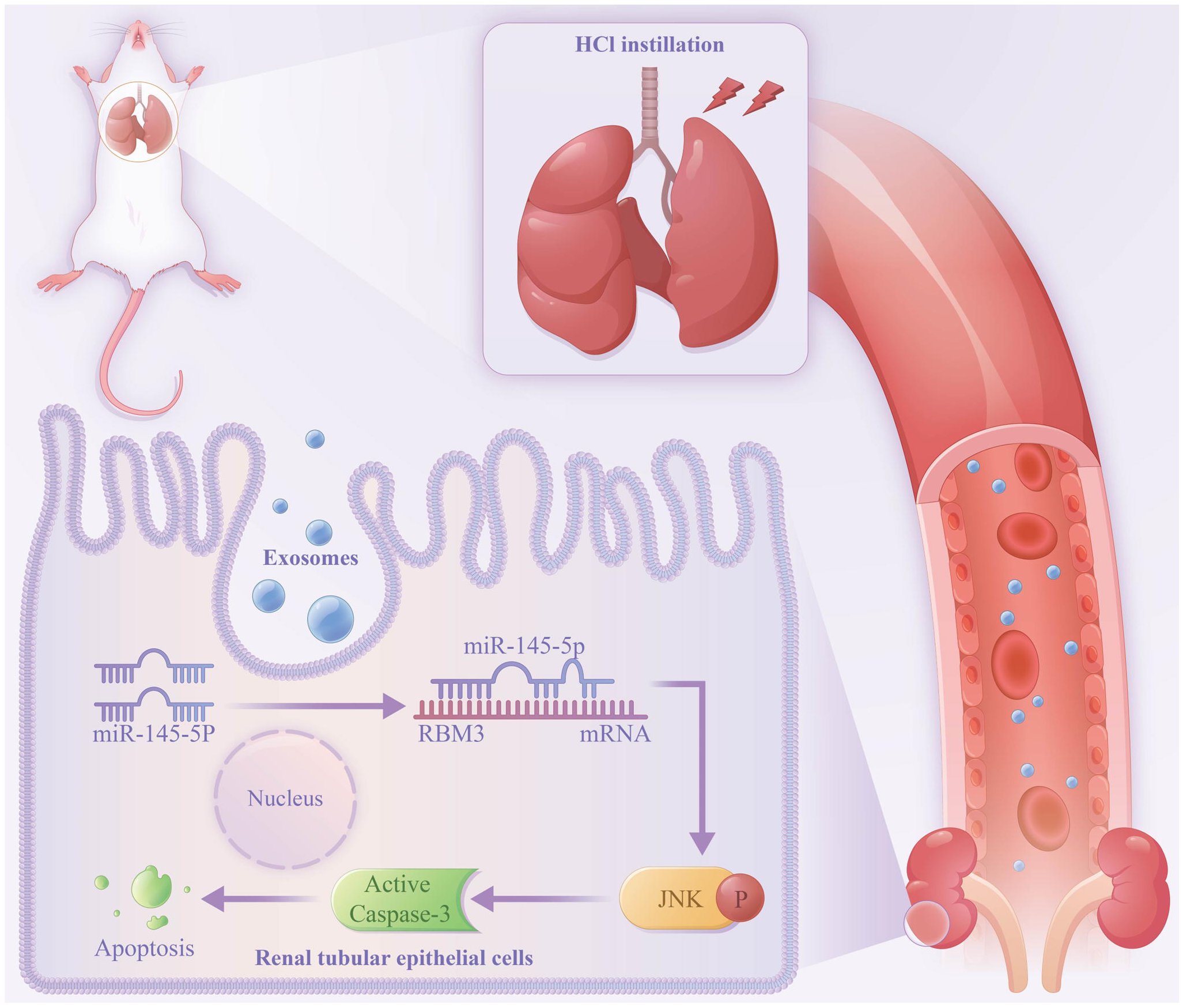

## REFERENCES

1. Bellani, G., Laffey, J.G., Pham, T., Fan, E., Brochard, L., Esteban, A., Gattinoni, L., van Haren, F., Larsson, A., McAuley, D.F., et al. (2016). Epidemiology, Patterns of Care, and Mortality for Patients With Acute Respiratory Distress Syndrome in Intensive Care Units in 50 Countries. Jama 315, 788–800. 10.1001/jama.2016.0291.

2. Panitchote, A., Mehkri, O., Hastings, A., Hanane, T., Demirjian, S., Torbic, H., Mireles-Cabodevila, E., Krishnan, S., and Duggal, A. (2019). Factors associated with acute kidney injury in acute respiratory distress syndrome. Annals of intensive care 9, 74. 10.1186/s13613-019-0552-5.

3. Darmon, M., Clec’h, C., Adrie, C., Argaud, L., Allaouchiche, B., Azoulay, E., Bouadma, L., Garrouste-Orgeas, M., Haouache, H., Schwebel, C., et al. (2014). Acute respiratory distress syndrome and risk of AKI among critically ill patients. Clinical journal of the American Society of Nephrology : CJASN 9, 1347–1353. 10.2215/cjn.08300813.

4. Tomasi, A., Song, X., Gajic, O., and Kashani, K. (2023). Kidney and lung crosstalk during critical illness: large-scale cohort study. Journal of nephrology 36, 1037–1046. 10.1007/s40620-022-01558-9.

5. Joannidis, M., Forni, L.G., Klein, S.J., Honore, P.M., Kashani, K., Ostermann, M., Prowle, J., Bagshaw, S.M., Cantaluppi, V., Darmon, M., et al. (2020). Lung-kidney interactions in critically ill patients: consensus report of the Acute Disease Quality Initiative (ADQI) 21 Workgroup. Intensive care medicine 46, 654–672. 10.1007/s00134-019-05869-7.

6. Kuiper, J.W., Groeneveld, A.B., Slutsky, A.S., and Plötz, F.B. (2005). Mechanical ventilation and acute renal failure. Critical care medicine 33, 1408–1415. 10.1097/01.ccm.0000165808.30416.ef.

7. Husain-Syed, F., Slutsky, A.S., and Ronco, C. (2016). Lung-Kidney Cross-Talk in the Critically Ill Patient. American journal of respiratory and critical care medicine 194, 402–414. 10.1164/rccm.201602-0420CP.

8. Liu, K.D., Glidden, D.V., Eisner, M.D., Parsons, P.E., Ware, L.B., Wheeler, A., Korpak, A., Thompson, B.T., Chertow, G.M., and Matthay, M.A. (2007). Predictive and pathogenetic value of plasma biomarkers for acute kidney injury in patients with acute lung injury. Critical care medicine 35, 2755–2761.

9. Kellum, J.A., and Prowle, J.R. (2018). Paradigms of acute kidney injury in the intensive care setting. Nature reviews. Nephrology 14, 217–230. 10.1038/nrneph.2017.184.

10. Pritchard, C.C., Cheng, H.H., and Tewari, M. (2012). MicroRNA profiling: approaches and considerations. Nature reviews. Genetics 13, 358–369. 10.1038/nrg3198.

11. Kalluri, R., and LeBleu, V.S. (2020). The biology, function, and biomedical applications of exosomes. Science (New York, N.Y.) 367. 10.1126/science.aau6977.

12. Lee, H., Zhang, D., Laskin, D.L., and Jin, Y. (2018). Functional Evidence of Pulmonary Extracellular Vesicles in Infectious and Noninfectious Lung Inflammation. Journal of immunology (Baltimore, Md. : 1950) 201, 1500–1509. 10.4049/jimmunol.1800264.

13. Parzibut, G., Henket, M., Moermans, C., Struman, I., Louis, E., Malaise, M., Louis, R., Misset, B., Njock, M.S., and Guiot, J. (2021). A Blood Exosomal miRNA Signature in Acute Respiratory Distress Syndrome. Frontiers in molecular biosciences 8, 640042. 10.3389/fmolb.2021.640042.

14. Wu, X., Wu, C., Gu, W., Ji, H., and Zhu, L. (2019). Serum Exosomal MicroRNAs Predict Acute Respiratory Distress Syndrome Events in Patients with Severe Community-Acquired Pneumonia. BioMed research international 2019, 3612020. 10.1155/2019/3612020.

15. Xu, F., Yuan, J., Tian, S., Chen, Y., and Zhou, F. (2020). MicroRNA-92a serves as a risk factor in sepsis-induced ARDS and regulates apoptosis and cell migration in lipopolysaccharide-induced HPMEC and A549 cell injury. Life sciences 256, 117957. 10.1016/j.lfs.2020.117957.

16. Linkermann, A., Chen, G., Dong, G., Kunzendorf, U., Krautwald, S., and Dong, Z. (2014). Regulated cell death in AKI. Journal of the American Society of Nephrology : JASN 25, 2689–2701. 10.1681/asn.2014030262.

17. Havasi, A., and Borkan, S.C. (2011). Apoptosis and acute kidney injury. Kidney international 80, 29–40. 10.1038/ki.2011.120.

18. Kulkarni, H.S., Lee, J.S., Bastarache, J.A., Kuebler, W.M., Downey, G.P., Albaiceta, G.M., Altemeier, W.A., Artigas, A., Bates, J.H.T., Calfee, C.S., et al. (2022). Update on the Features and Measurements of Experimental Acute Lung Injury in Animals: An Official American Thoracic Society Workshop Report. American journal of respiratory cell and molecular biology 66, e1–e14. 10.1165/rcmb.2021-0531ST.

19. Matute-Bello, G., Downey, G., Moore, B.B., Groshong, S.D., Matthay, M.A., Slutsky, A.S., and Kuebler, W.M. (2011). An official American Thoracic Society workshop report: features and measurements of experimental acute lung injury in animals. American journal of respiratory cell and molecular biology 44, 725–738. 10.1165/rcmb.2009-0210ST.

20. Wang, F., Flanagan, J., Su, N., Wang, L.C., Bui, S., Nielson, A., Wu, X., Vo, H.T., Ma, X.J., and Luo, Y. (2012). RNAscope: a novel in situ RNA analysis platform for formalin-fixed, paraffin-embedded tissues. The Journal of molecular diagnostics : JMD 14, 22–29. 10.1016/j.jmoldx.2011.08.002.

21. Caruso, P., Dempsie, Y., Stevens, H.C., McDonald, R.A., Long, L., Lu, R., White, K., Mair, K.M., McClure, J.D., Southwood, M., et al. (2012). A role for miR-145 in pulmonary arterial hypertension: evidence from mouse models and patient samples. Circulation research 111, 290–300. 10.1161/circresaha.112.267591.

22. Yue, J., and López, J.M. (2020). Understanding MAPK Signaling Pathways in Apoptosis. International journal of molecular sciences 21. 10.3390/ijms21072346.

23. Liu, J., and Lin, A. (2005). Role of JNK activation in apoptosis: a double-edged sword. Cell research 15, 36–42. 10.1038/sj.cr.7290262.

24. Min, W., Lin, Y., Tang, S., Yu, L., Zhang, H., Wan, T., Luhn, T., Fu, H., and Chen, H. (2008). AIP1 recruits phosphatase PP2A to ASK1 in tumor necrosis factor-induced ASK1-JNK activation. Circulation research 102, 840–848. 10.1161/circresaha.107.168153.

25. Zhang, H., Zhang, H., Lin, Y., Li, J., Pober, J.S., and Min, W. (2007). RIP1-mediated AIP1 phosphorylation at a 14-3-3-binding site is critical for tumor necrosis factor-induced ASK1-JNK/p38 activation. The Journal of biological chemistry 282, 14788–14796. 10.1074/jbc.M701148200.

26. Tang, X., Yu, Q., Wen, X., Qi, D., Peng, J., He, J., Deng, W., Zhu, T., Zhao, Y., and Wang, D. (2020). Circulating Exosomes From Lipopolysaccharide-Induced Ards Mice Trigger Endoplasmic Reticulum Stress in Lung Tissue. Shock (Augusta, Ga.) 54, 110–118. 10.1097/shk.0000000000001397.

27. Jiang, K., Yang, J., Guo, S., Zhao, G., Wu, H., and Deng, G. (2019). Peripheral Circulating Exosome-Mediated Delivery of miR-155 as a Novel Mechanism for Acute Lung Inflammation. Molecular therapy : the journal of the American Society of Gene Therapy 27, 1758–1771. 10.1016/j.ymthe.2019.07.003.

28. Pan, T., Jia, P., Chen, N., Fang, Y., Liang, Y., Guo, M., and Ding, X. (2019). Delayed Remote Ischemic Preconditioning ConfersRenoprotection against Septic Acute Kidney Injury via Exosomal miR-21. Theranostics 9, 405–423. 10.7150/thno.29832.

29. Sancho-Martínez, S.M., López-Novoa, J.M., and López-Hernández, F.J. (2015). Pathophysiological role of different tubular epithelial cell death modes in acute kidney injury. Clinical kidney journal 8, 548–559. 10.1093/ckj/sfv069.

30. Ren, Q., Guo, F., Tao, S., Huang, R., Ma, L., and Fu, P. (2020). Flavonoid fisetin alleviates kidney inflammation and apoptosis via inhibiting Src-mediated NF-κB p65 and MAPK signaling pathways in septic AKI mice. Biomedicine & pharmacotherapy = Biomedecine & pharmacotherapie 122, 109772. 10.1016/j.biopha.2019.109772.

31. Medrano, S., Sequeira-Lopez, M.L., and Gomez, R.A. (2014). Deletion of the miR-143/145 cluster leads to hydronephrosis in mice. The American journal of pathology 184, 3226–3238. 10.1016/j.ajpath.2014.08.012.

32. Deng, L., Blanco, F.J., Stevens, H., Lu, R., Caudrillier, A., McBride, M., McClure, J.D., Grant, J., Thomas, M., Frid, M., et al. (2015). MicroRNA-143 Activation Regulates Smooth Muscle and Endothelial Cell Crosstalk in Pulmonary Arterial Hypertension. Circulation research 117, 870–883. 10.1161/circresaha.115.306806.

33. Wellmann, S., Truss, M., Bruder, E., Tornillo, L., Zelmer, A., Seeger, K., and Bührer, C. (2010). The RNA-binding protein RBM3 is required for cell proliferation and protects against serum deprivation-induced cell death. Pediatric research 67, 35–41. 10.1203/PDR.0b013e3181c13326.

34. Ma, R., Zhao, L.N., Yang, H., Wang, Y.F., Hu, J., Zang, J., Mao, J.G., Xiao, J.J., and Shi, M. (2018). RNA binding motif protein 3 (RBM3) drives radioresistance in nasopharyngeal carcinoma by reducing apoptosis via the PI3K/AKT/Bcl-2 signaling pathway. American journal of translational research 10, 4130–4140.

35. Feng, J., Pan, W., Yang, X., Long, F., Zhou, J., Liao, Y., and Wang, M. (2021). RBM3 Increases Cell Survival but Disrupts Tight Junction of Microvascular Endothelial Cells in Acute Lung Injury. The Journal of surgical research 261, 226–235. 10.1016/j.jss.2020.12.041.

