## Supplementary figures and images for "Exosomal miR-145-5p promotes apoptosis of renal tubule epithelial cells through the JNK signalling pathway"

### Supplemental figure 1

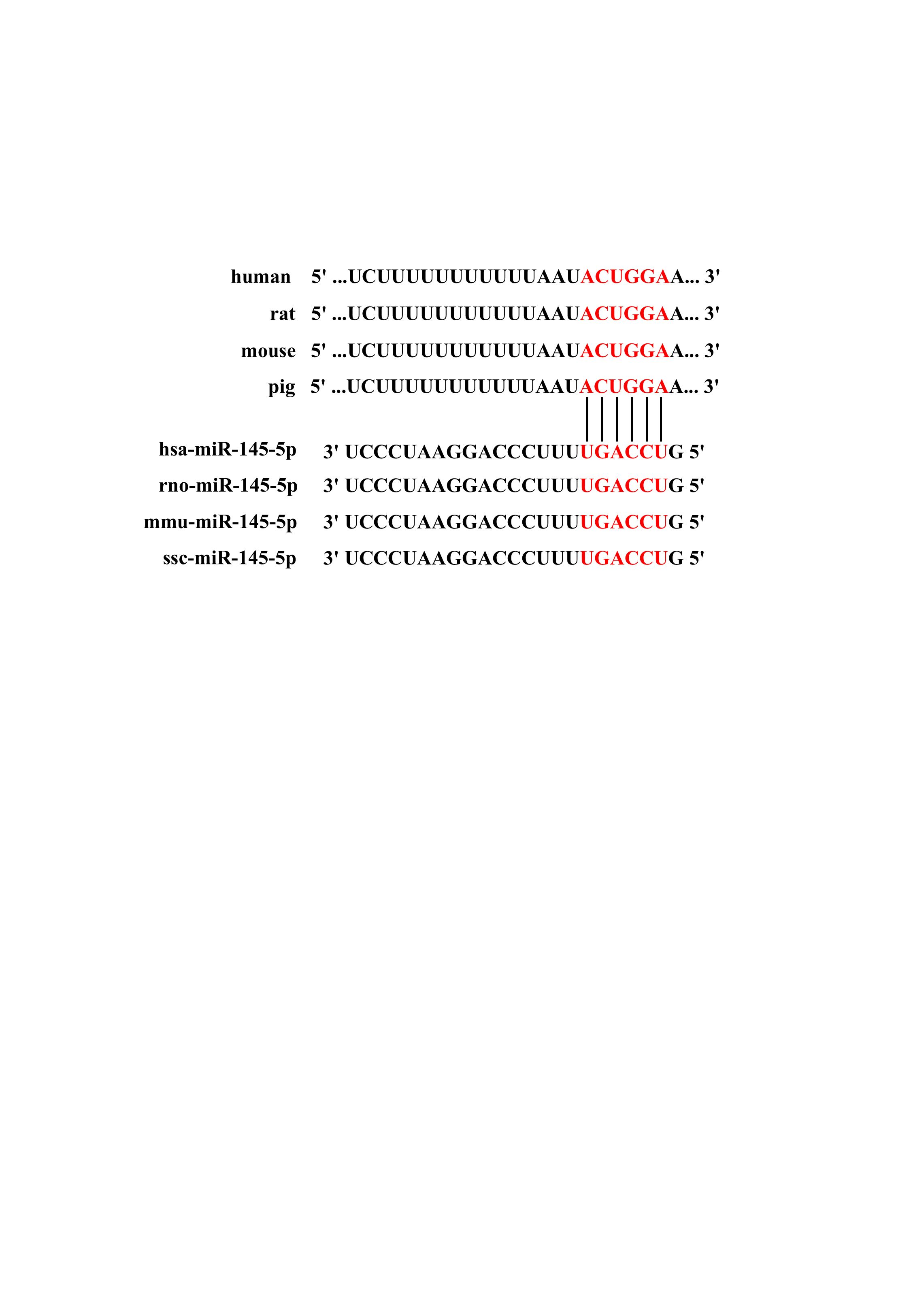

### Supplementary figure 2

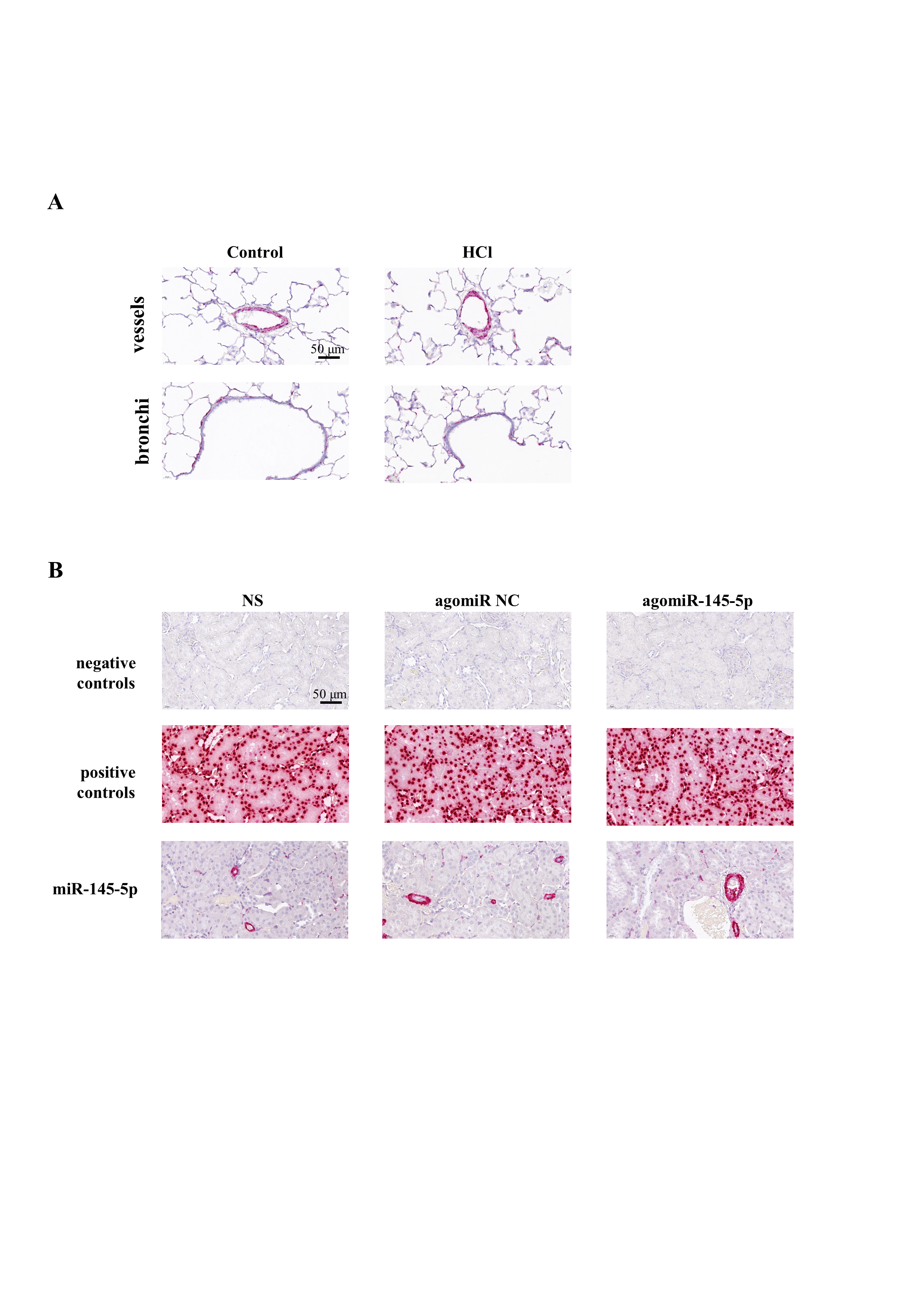
